# Arsinothricin Biosynthesis Involving a Noncanonical Radical SAM Enzyme for C-As Bond Formation

**DOI:** 10.1101/2024.02.01.577332

**Authors:** Yadi Yao, Jiale He, Min Dong

## Abstract

Arsinothricin is a potent antibiotic secreted by soil bacteria. The biosynthesis of Arsinothricin was proposed to involve two steps. The first step is C-As bond formation between trivalent As and the 3-amino-3-carboxypropyl (ACP) group of S-adenosyl-L-methionine (SAM), which is catalyzed by the protein ArsL. However, the reaction has not been verified in vitro, and ArsL has not been characterized in detail. Interestingly, ArsL contains a CxxxCxxC motif and thus belongs to the radical SAM enzyme superfamily, the members of which cleave SAM and generate a 5′-deoxyadenosyl radical. Here, we found that ArsL cleaves the C_γ,Met_–S bond of SAM and generates an ACP radical that resembles Dph2, a noncanonical radical SAM enzyme involved in diphthamid biosynthesis. As Dph2 does not contain the CxxxCxxC motif, ArsL is a unique noncanonical radical SAM enzyme that contains this motif but generates an ACP radical. Together with the methyltransferase ArsM, we successfully reconstituted arsinothricin biosynthesis in vitro. ArsL has a conserved RCCLKC motif in the C-terminal sequence and belongs to the RCCLKC-tail radical SAM protein subfamily. By truncation, we showed that this motif binds to the substrate arsenite and is highly important for its activity. Our results suggested that ArsL is a noncanonical radical SAM enzyme with a canonical radical SAM enzyme motif, implying that more noncanonical radical SAM chemistry may exist within the radical SAM enzyme superfamily.

**TOC:** 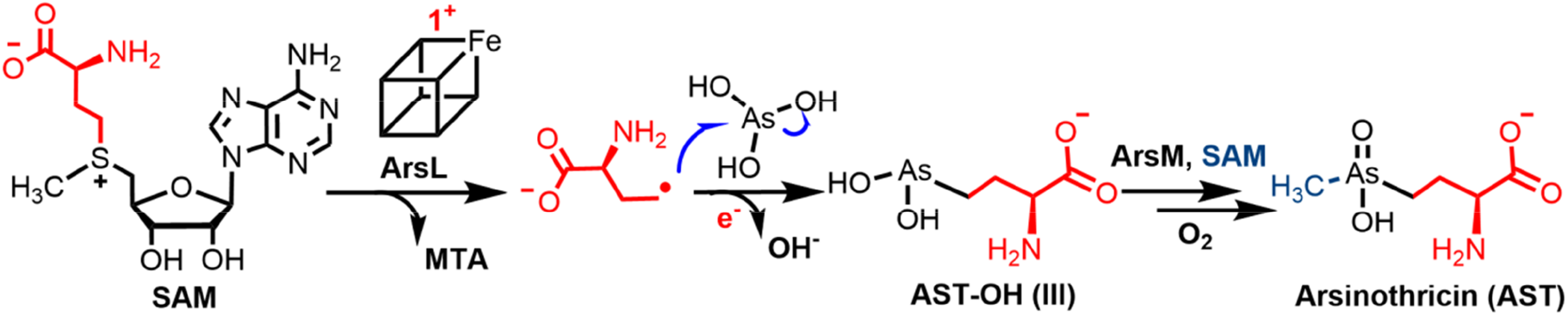

---

Arsenic is a widely distributed toxic metalloid in nature and is found mainly as inorganic arsenic species, such as arsenite (III) and arsenate (V), which are toxic contaminants in the environment.^1^ Despite its toxicity, inorganic arsenic has been successfully used to treat a variety of human diseases^2^, such as acute promyelocytic leukemia (APL)^3^ and multiple myeloma^4^. As a detoxic strategy,^5,6^ bacteria convert inorganic As to organic compounds, such as methylarsenic, dimethyl arsenic, trimethyl arsenic, arsenosugars^7^ and its derivatives arsenobetaine^8^ and arsenolipids.^9,10^ Recently, researchers discovered a natural arsenic compound arsinothricin (AST), which is produced by soil bacteria in the rice rhizosphere as a defensive compound.^11^ Arsinothricin is a potent inhibitor of glutamine synthetase and a promising antibiotic for various pathogenic microorganisms, including drug-resistant species.^12,13^ Therefore, arsinothricin is a promising drug candidate and its efficient preparation and further study are highly desirable. The biosynthesis of Arsinothricin was proposed to involve two steps.^14^ The first step is the transfer of the 3-amino-3-carboxyl propal (ACP) group from S-adenosinmethionine (SAM) to arenite, generating the trivalent AST-OH. The second step is the methylation of AST-OH by the SAM-dependent methyltransferase ArsM to produce trivalent arsinothricin, which is readily oxidized by air to form arsinothricin (Figure 1). The second step of arsinothricin biosynthesis has been studied both in vivo^15^ and with chemically synthesized trivalent AST-OH and purified ArsM in vitro.^16^ However, the intriguing first step in the biosynthesis route for C-As bond formation with ArsL catalysis remains unclear.

**Figure 1.**
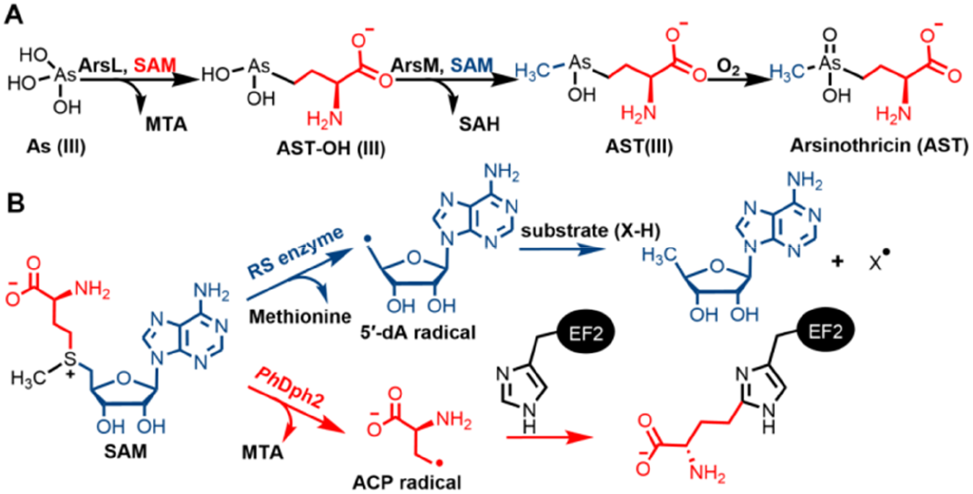
(A) Proposed arsinothricin biosynthesis pathway. (B) The radical SAM (RS) enzyme and noncanonical radical SAM enzyme Dph2 catalyzed SAM cleavage reactions; to simplify the scheme, organometallic intermediates that function as stable radicals are not shown.

As the protein sequence of ArsL contains a CxxxCxxC motif, the enzyme was predicted to be a member of the radical SAM enzyme superfamily, which is the largest protein family with more than 70,000 members.^17–19^ The radical SAM enzyme binds to the 4Fe-4S cluster via the conserved CxxxCxxC motif.^20,21^ The reduced 4Fe-4S cluster cleaves SAM to generate the 5′-deoxyadenosyl (5′-dA) radical, which mostly absorbs a hydride from the substrate to produce a substrate radical;^21^ this radical catalyzes numerous challenging reactions or directly adds to the substrate in several cases.^22,23^ Interestingly, the proposed activity of ArsL in arsinothricin biosynthesis involves the transfer of the ACP group from SAM to arsenite.^14^ This unusual SAM cleavage resembles the noncanocial radical SAM enzyme Dph2 in diphthamide biosynthesis.^24–26^ Dph2 cleaves SAM and generates a 4Fe-4S cluster-stabilized ACP radical to modify the histidine residue of the substrate protein EF2 (Figure 1). Interestingly, the special SAM cleavage activity of Dph2 resulted from the distinct geometry of SAM binding with 4Fe-4S, which was conjugated by three cysteines from different domains of the protein instead of the CxxxCxxC motif.^25^ The Broderick group^27^ and our group^28^ recently demonstrated that glycerol dehydrogenase-activating enzyme GD-AE, the only previously assumed noncanonical radical SAM enzyme with the CxxxCxxC motif, is a canonical radical SAM enzyme that generates the 5′-dA radical. Moreover, many enzymes, such as TWY2^29^, Tsr3^30^ and AzeJ^31^, cleave the C_γ,Met_-S bond of SAM by a nucleophilic mechanism.^32^ To date, no radical SAM enzymes with the CxxxCxxC motif have been reported to cleave the C_γ,Met_-S bond of SAM and generate the ACP radical, as does Dph2. In terms of the CxxxCxxC motif in ArsL, it remains unknown whether ArsL is a radical SAM enzyme or only binds a 4Fe-4S cluster but catalyzes a nucleophilic reaction, such as the SAM-dependent methyltransferase TsrM^33,34^ in the biosynthesis of thiostrepton.

The *Burkholderia gladioli* ArsL protein was overexpressed in *Escherichia coli* and purified under anaerobic conditions (Figure S1). The purified ArsL had a brown color and UV absorption at 410 nm, which disappeared when dithionite was added as the reductant, typical for the [4Fe-4S]^2+^ cluster of radical SAM enzymes (Figure S2). Analyses of the iron and sulfur contents of ArsL showed 1.2±0.1 and 2.3±0.4 equiv. of iron and sulfur, respectively, per protein, suggesting the presence of 0.3 [4Fe-4S]^2+^ cluster loading. We first tested the activity of ArsL to determine whether it could cleave SAM to generate 5′-dA, MTA, or both products as 5′-dA and SAH detected in the reaction of radical SAM methyltransferases.^35^ Dithionite was used as the reductant in the reaction. High-performance liquid chromatography (HPLC) showed that in the reaction containing ArsL, SAM and dithionite, SAM was completely converted to MTA, which was further confirmed by high-resolution mass spectrometry (HR-MS) (Figure 2). No MTA was formed in the control reaction without dithionite, indicating that the SAM cleavage activity depends on the reduced [4Fe-4S]^1+^ cluster. In the control reaction without ArsL, only trace amounts of MTA were detected due to SAM decomposition (Figure 2). This result suggested that ArsL is different from the conventional radical SAM enzyme, which cleaves SAM to generate 5′-dA. The dependence on reductant suggested that ArsL did not catalyze SAM cleavage by a nucleophilic mechanism like TsrM^33,34^ but rather through a radical mechanism like Dph2^24–26^. To confirm the radical mechanism of ArsL, we detected the dithionite radical-quenched ACP product, as has been detected for Dph2 and the quenched product of the 5′-dA radical in many conventional radical SAM enzymes.^36,37^ Homocysteine sulphinic acid (HSA) was detected in the reaction by LC-MS but was not detected in the control reactions without dithionite. The result was further validated by derivatizing the reaction product with 2,4-dinitrofluorobenzene (DNFB) and analysis by LC-HRMS. Derivatized HSA was detected only in the reaction and not in the controls without dithionite or ArsL (Figure 3).

**Figure 2.**
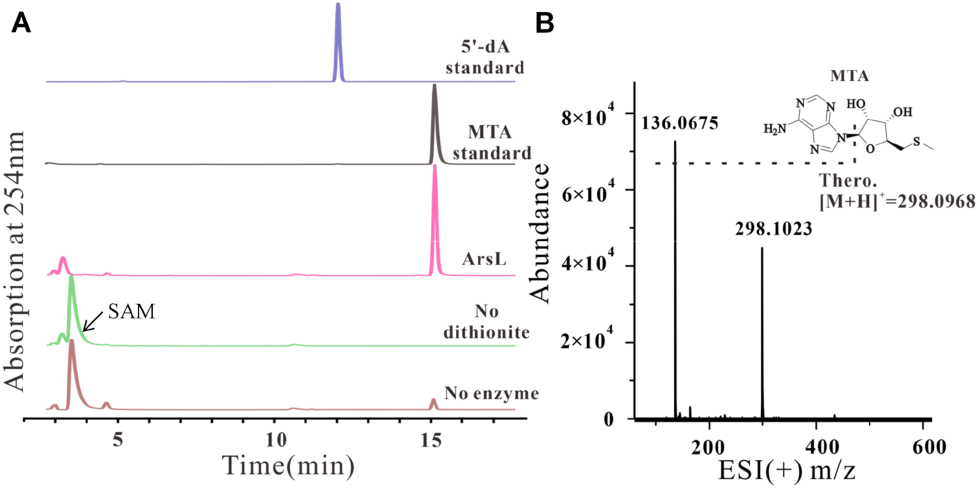
SAM cleavage reaction of ArsL. Left: HPLC traces of the ArsL reactions. The full reaction mixture contained ArsL, SAM and dithionite. SAM was not cleaved without ArsL or dithionite. Right: HR-MS of the isolated MTA peak in the full reaction.

**Figure 3.**
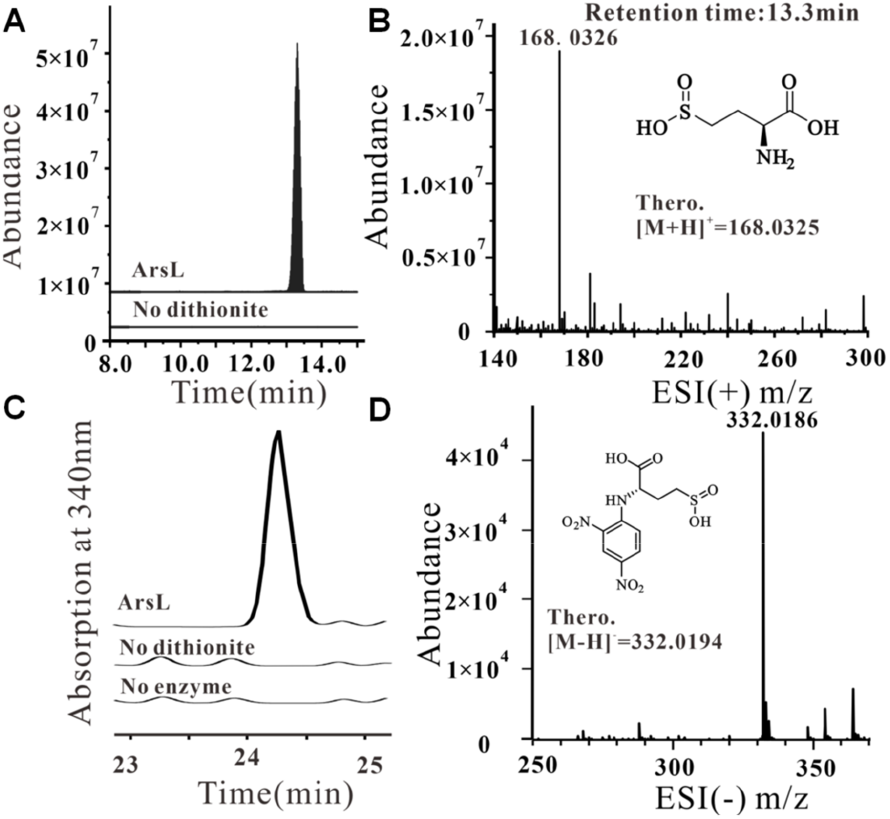
Detection of the dithionite-quenched ACP radical product HSA. A) Extracted ion chromatographs monitoring the formation of HSA in the reaction. B) Positive ionization high-resolution mass spectrum of the HSA peak eluted at 13.3 min. C) Extracted ion chromatographs monitoring the formation of the DNFB derivative from the HSA product. No product formed in the negative controls without dithionite or ArsL. D) Negative ionization high-resolution mass spectrum obtained for the DNFB derivative of HSA.

We next investigated the ArsL reaction with the substrate arsenite (As(III)). Due to the structural similarity of dithionite with the substrate As(III) and the presence of the quenched ACP product HAS with dithionite radicals, we suspected that dithionite may compete with As(III). Therefore, in addition to dithionite, we also performed reactions with other small-molecule reductants used for radical SAM enzyme reduction, including methylviologen (MV)/NADPH and titanium citrate. HPLC showed that SAM was converted to MTA in all three reactions with different reductants, whereas no MTA was generated in the absence of a reductant or ArsL (Figure 4). Interestingly, more MTA was produced with dithiontie compared to the other two reductants. Marfey’s method was used to produce L-FDAA derivatives of the product.^38^ Hydrogen peroxide was subsequently used to oxidize As(III) and release the product from the protein. Derivatized AST-OH (V) was detected by HR-MS in all three reactions with different reductants. Interestingly, much less product was detected in the reaction with dithionite than in the reactions with MV/NADPH or titanium citrate. This result, together with more MTA detected in the dithionite reaction, suggested that dithionite competed with the substrate As(III) and induced the redudant SAM cleavage reaction. Therefore, we used titanium citrate or MV/NADPH as reductants in our subsequent experiments. To further confirm the product formation, we used HR-MS to directly detect AST-OH(III) without derivatization. AST-OH(III) was detected in the reaction and structurally confirmed by tandem MS, which was identical to the results from a previous report on AST-OH identification (Figure S3).^11^

**Figure 4.**
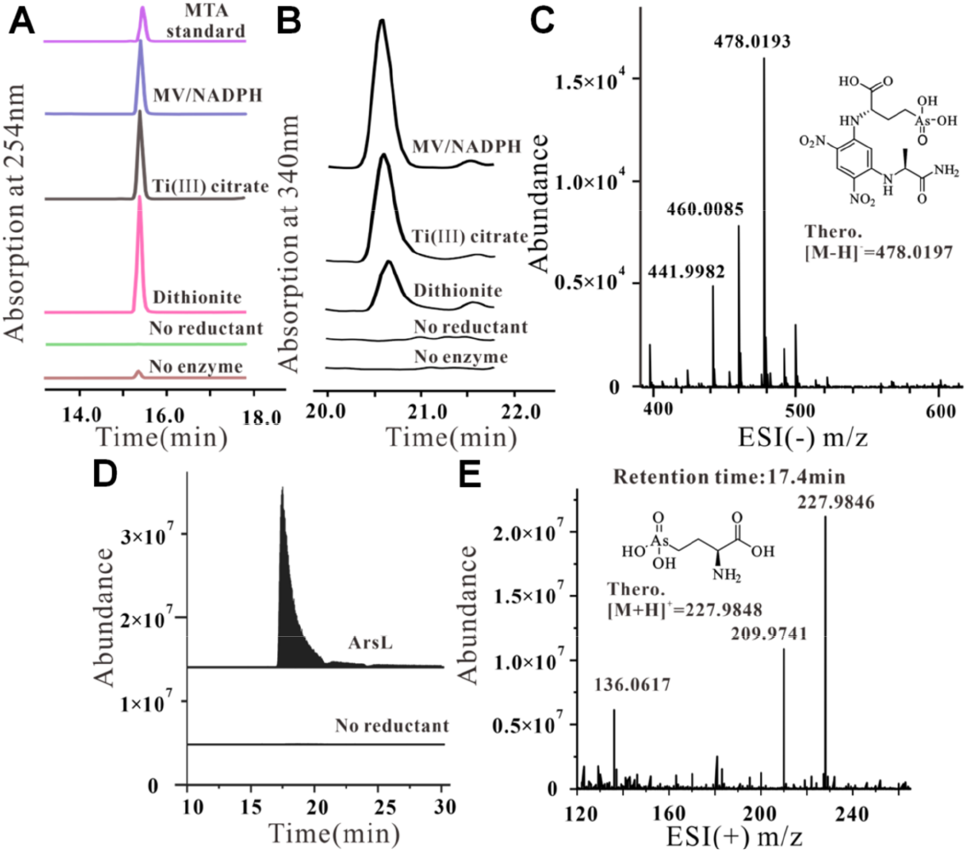
Detection of AST-OH from the ArsL reactions. A) HPLC traces of the SAM cleavage reactions by ArsL in the presence of As(III) with different reductants. B) HPLC elution profiles of the L-FDAA derivatives of AST-OH. C) Negative ionization high-resolution mass spectrum of the L-FDAA derivatives of AST-OH. D) Extracted ion chromatographs monitoring the formation of AST-OH in the ArsL reaction. No products formed in the negative controls without reductant. E) Positive ionization high-resolution mass spectrum of the AST-OH peak eluting at 17.4 min.

With the active ArsL, we then tested the tandem reaction between ArsL and *Bg*ArsM to reconstitute the biosynthesis of AST in vitro. The reaction mixture containing SAM, ArsL, ArsM, As(III) and titanium citrate was incubated at 28°; for 5 h. The reaction mixture without ArsM was used as a control for the cascade reaction. HPLC showed that in the reaction with ArsL and ArsM, SAM was converted to MTA and SAH, suggesting that ACP and methyl transfer both occurred (Figure 5). In the control reaction without ArsM, only MTA was produced. These two reactions were then treated with H_2_O_2_ to oxidize and release the products, which were subsequently derivatized with L-FDAA to facilitate product detection. The L-FDAA derivative of AST was detected in the cascade reaction, together with the L-FDAA derivative of AST-OH. No AST was detected, but only AST-OH was detected in the ArsL reaction. To further confirm that AST formed, we detected the product directly without derivatization. AST was detected by HR-MS only in the cascade reaction. The tandem mass spectrum of AST, which was identical to that in the literature,^11^ verified the formation of the product (Figure S4).

**Figure 5.**
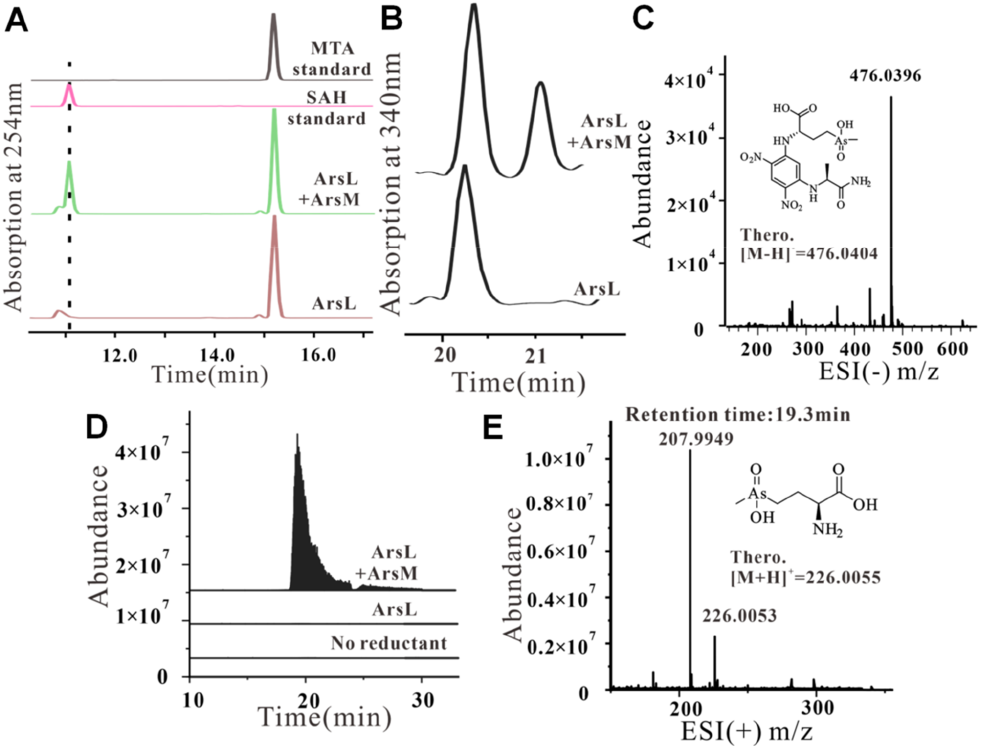
Detection of AST from the cascade reaction of ArsL with ArsM. A) HPLC traces of the SAM cleavage reactions showing that both MTA and SAH were produced in the cascade reaction involving ArsL and ArsM. Only MTA was generated in the ArsL reaction. B) HPLC elution profiles of the L-FDAA derivatives of AST. No products formed in the ArsL reaction. C) Negative ionization high-resolution mass spectrum of the L-FDAA derivatives of AST. D) Extracted ion chromatographs monitoring the formation of AST in the cascade reaction of ArsL with ArsM. No AST formed in the ArsL reaction. E) Positive ionization high-resolution mass spectrum of the AST-OH peak eluting at 19.3 min.

To further study the biosynthesis of AST, we performed a stepwise cascade reaction with ArsL and ArsM. Two identical reactions involving ArsL, SAM, As(III) and titanium citrate were performed first. After the mixture was incubated for 4 h, one reaction was directly applied to ArsM for methyltransfer (reaction 1). Another reaction was oxided by H_2_O_2_, after which the mixture was lyophilized to remove additional H_2_O_2_ and added to ArsM (reaction 2). After reaction 2 was incubated for 8 h, these two reactions were derivatized with L-FDAA for product detection. HPLC and MS showed that AST-OH and AST were produced in reaction 1 without oxidation after the ArsL reaction. Reaction 2 produced only AST-OH (Figure S6). This finding suggested that the product of ArsL was AST-OH(III), which was the substrate of ArsM. After treatment with H_2_O_2_, the ArsL product was oxidized to AST-OH(IV), which was not a substrate of ArsM. A previous study on ArsM with synthesized AST-OH(IV) showed that ArsM methylated chemically reduced AST-OH(III).^16^. Our results were consistent with these findings and provided direct evidence that the product of ArsL was AST-OH(III).

Sequence alignment of ArsL proteins from different species revealed a conserved RCCLKC motif at the C-terminal end of the sequence (Figure S7). Many radical SAM enzymes with extra cysteine motifs in the sequence bind to auxiliary 4Fe-4S clusters.^39^ To investigate the function of the three conserved cysteines at the C-terminal end of ArsL, we first prepared truncated ArsL without the RCCLKC motif as ArsL_ΔC_. After iron-sulfur cluster reconstitution and iron and sulfur content analyses, ArsL_ΔC_ was found to contain similar amounts of iron and sulfur as the wild-type protein. This result suggested that the RCCLKC motif did not bind to an auxiliary [4Fe-4S]^2+^ cluster. Studies on ArsM have shown that ArsM contains a conserved four-cysteine motif for As(III) binding, which facilitates the nucleophilic substitution of As for methylation. Given that ArsL also uses As(III) as a substrate, we speculated that the RCCLKC motif could also serve as a binding site for As(III).

To test the function of the RCCLKC motif of ArsL, we first validated the SAM cleavage activity of ArsL_ΔC_ without the substrate As(III). Like wild-type ArsL, ArsL_ΔC_ efficiently cleaved SAM to MTA (Figure 6a). Derivatization with L-FDAA, HPLC and HR-MS revealed that the wild-type ArsL reaction and the ArsL_ΔC_ reaction generated 2-aminobutyric acid (ABA), the product of the ACP radical quenched by a proton (Figure 6b, 6c). These results suggested that the RCCLKC motif does not affect the SAM cleavage activity of ArsL without the substrate As(III). When the substrate As(III) was present, ArsL produced AST-OH and a small amount of ABA. However, ArsL_ΔC_ only generated ABA and not AST-OH. These results suggested that the RCCLKC motif binds to the substrate As(III). Therefore, the function of the RCCLKC motif in ArsL was clearly established. In addition to As(III), we also tested phosphorous acid and two other analogues to ArsL and ArsL_ΔC_, no desired products were detected but only ABA (Figure S8). Apparently, these phosphorus analogues of As(III) were not substrates of ArsL.

**Figure 6.**
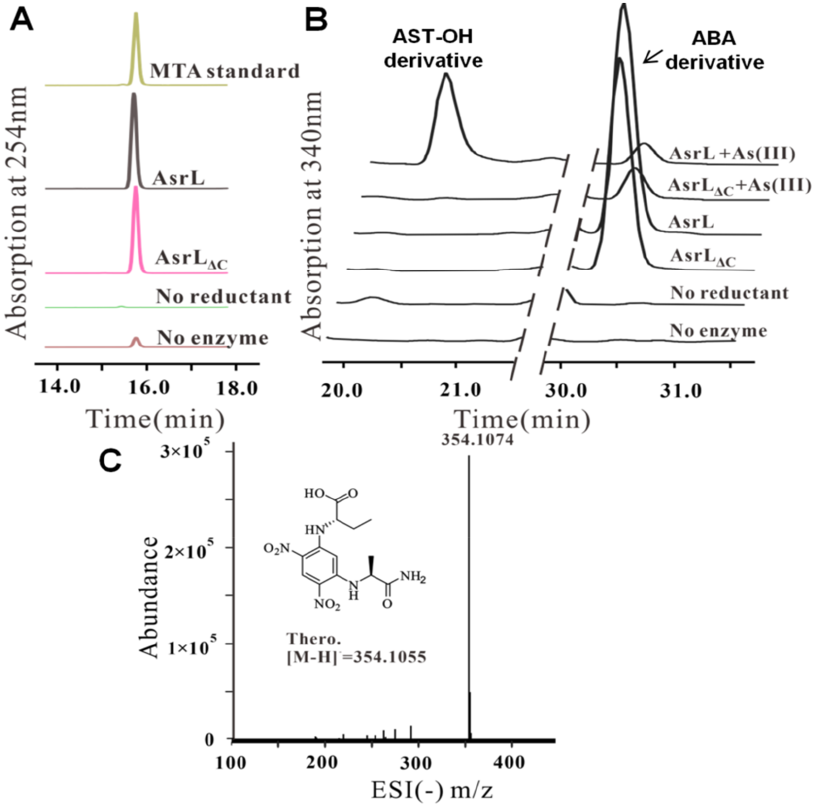
Functional study of the RCCLKC motif of ArsL. A) HPLC traces of the SAM cleavage reactions showing that the wild-type ArsL and RCCLKC motif-truncated ArsLΔC can cleave SAM to MTA. B) HPLC elution profiles for the L-FDAA derivatives of AST-OH in the ArsL wildtype reaction and L-FDAA derivatives of ABA in the ArsLΔC reaction. C) Negative ionization high-resolution mass spectrum of the L-FDAA derivatives of ABA.

In the NCBIfam database, a subfamily of the radical SAM enzyme superfamily was named the RCCLKC-tail radical SAM protein (NCBI HMM accession NF040542.1). This subfamily included 147 sequences, 39 of which were annotated as the arsinothricin biosynthesis radical SAM protein ArsL. The other 108 sequences were listed as RCCLKC-tail radical SAM proteins with unknown function (Figure S9). Our study on the activity of ArsL, especially the function of the RCCLKC motif in As binding, suggested that these 108 sequences could all encode noncanonical radical SAM enzymes as ArsL that trasfer the ACP group to As(III).

In summary, we demonstrated that ArsL is a noncanonical radical SAM enzyme that transfers the ACP group from SAM to arsenic acid by a radical mechanism for C-As bond formation in arsinothricin biosynthesis. For the first time, we purified ArsL and identified the reaction products (Figure 7). We further reconstituted the biosynthesis of Arsinothricin in vitro with the methyltransferase ArsM. All three C-S bonds of SAM were used for the biosynthesis of arsenic derivatives in nature, as demonstrated by the nucleophilic mechanism that underlies C_methyl_-S bond cleavage of ArsM, the radical mechanism that underlies C_ad_-S bond cleavage of ArsS in arsenosugar biosynthesis^22^, and the radical mechanism that underlies C_γ,Met_-S bond cleavage of ArsL. More importantly, ArsL is the second noncanonical radical SAM enzyme that catalyzes the ACP transfer reaction through a radical mechanism but is the only noncanonical radical SAM enzyme with the CXXXCXXC motif. This motif is the characteristic sequence for the radical SAM enzyme and was used to identify over 700,000 members of the radial SAM enzyme superfamily. Based on our research on ArsL, these members could contain additional noncanonical radical SAM enzymes with stunning and interesting chemistry.

**Figure 7.**
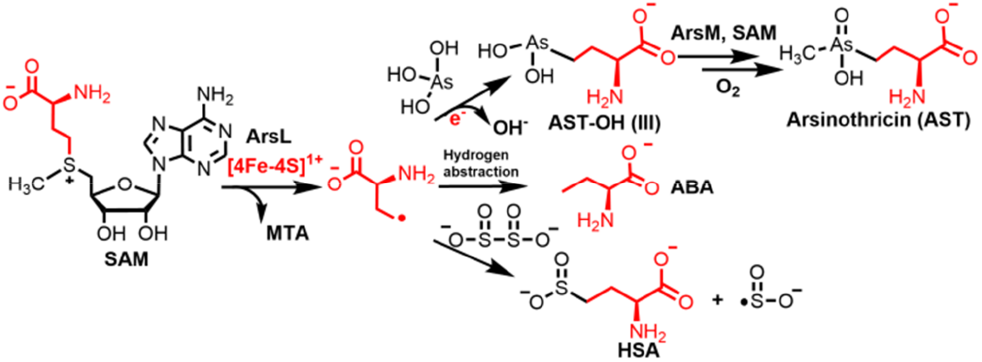
The proposed reaction mechanism of ArsL

## ASSOCIATED CONTENT

The supporting information contains the experimental materials and methods and supporting figures. This material is available free of charge on the ACS Publications website.

## AUTHOR INFORMATION

## ACKNOWLEDGMENT

This work is supported by the National Natural Science Foundation of China (22277088, 21977079), the Natural Science Foundation of Tianjin, China (20JCYBJC01100), the State Key Laboratory of Bio-organic and Natural Products Chemistry, CAS (SKLBNPC201108) and the Key-Area Research and Development Program of Guangdong Province (2020B0303070002).

