## Supplemental Information for "Arsinothricin Biosynthesis Involving a Noncanonical Radical SAM Enzyme for C-As Bond Formation"

**Materials and methods**

**A. Reagents and Methods** **S2-S7**
**B. Supplementary Tables and Figures** **S8-S15**
**C. References** **S16**

**Reagents and Methods:**

**Reagents.**

SAM was purchased from SolarBio (Shanghai, China). Arsenious acid (As (III)) was purchased from TMRM (Jiangsu, China). Dimethylphosphinous acid ($C_2H_7OP$) was purchased from Bidepharm (Shanghai, China). Trimethylphosphorus ($C_3H_9P$) was purchased from TCI (Shanghai, China). The pTrc-HisA (Kana) vector was constructed from pTrc-HisA (Amp) by transforming the antibiotic resistance from ampicillin to kanamycin. The *E. coli* BL21 Δmtn (DE3) strain was generated by knocking out SAH nucleosidase from the *E. coli* BL21 (DE3) strain.

**Cloning and expression of ArsL.**

The codon-optimized gene fragment of ArsL (accession number WP_219608243) from *Burkholderia gladioli* GSRB05 (NZ_JAGSIB010000059.1)[1] was synthesized by GENE CREATE (Wuhan, China), and inserted into the vector pTrc-HisA (Amp). To enhance the binding affinity between ArsL and Ni-NTA resin, a flexible protein linker consisting of 6×glycine was added between the His tag and ArsL via Gibson assembly. Using the Minerva Super Fusion Cloning Kit (UE EVERBRIGHT), the gene with 6×glycine was cloned and inserted into the vector pTrc-HisA (Kana). The primers used are listed in Table S1. The plasmid was subsequently used to transform the *E. coli* Rosetta (DE3) strain. The cells were grown in 1 liter LB medium with 30 μg/mL kanamycin and 20 μg/mL chloramphenicol at 37 °C and 180 rpm. When the $OD_{600}$ reached 0.8, the cultures were cooled down to 16 °C and supplemented with $FeCl_3$, Fe $(NH_4)_2(SO_4)_2$ and L-cysteine to final concentrations of 50 μM, 50 μM and 400 μM, respectively. Protein expression was induced by 1mM isopropyl-β-D-thiogalactopyranoside (IPTG), after which the culture flasks were sealed to limit the amount of oxygen in the system. The cells were incubated in a shaker at 16 °C and 180 rpm for 18-20 h before being harvested.

**Construction of the truncation of ArsL without the CCLKC motif.**

The C-terminal truncation of ArsL without the CCLKC motif ($ArsL_{\Delta C}$) was constructed by Gibson assembly using a Minerva Super Fusion Cloning Kit (UE EVERBRIGHT). The primers used are listed in Table S1.

**Purification of ArsL.**

Protein purification was performed in an anaerobic chamber, or protein purified under aerobic condition and [4Fe-4S] was reconstituted in an anaerobic chamber, as described below. The cell pellets from 2 liters culture were suspended in 40 mL lysis buffer (300 mM NaCl, 5 mM imidazole, and 50 mM $NaH_2PO_4$, 10% (v/v) glycerol at pH 7.8). Then, 1 mM PMSF, 120 mg lysozyme, 2 mg DNase I and 2 mM thioglycol were added, and the cell suspension was incubated at RT for 15 min and frozen in a -80°C freezer. The mixture was thawed and disrupted by sonication (Scientz, China) at 0 °C for 30 min (power 300 W, working 4 sec with a 6 sec interval). Cell debris was removed by centrifugation at 14,000g (Cence, China) for 40 min. The supernatant was incubated for 1 h with 3 mL of Ni-NTA resin (GenScript) preequilibrated with the lysis buffer. The Ni-NTA resin was loaded onto a polypropylene column and washed with 40 mL lysis buffer, followed by 300 mL of 30 mM imidazole and 80mL of 50mM imidazole in lysis buffer. ArsL was eluted from the column with elution buffer (200 mM and 500 mM imidazole in lysis buffer). The elution fractions were buffer-exchanged to 300 mM NaCl, 50 mM $NaH_2PO_4$, and 10% (v/v) glycerol at pH 8.0 using a 10-DG desalting column (GE Healthcare). The purified proteins were concentrated using Amicon Ultra 15 centrifugal filter devices (Millipore) (Figure S1).

**In vitro Fe-S cluster reconstitution of ArsL.**

Aerobically purified ArsL proteins were degassed by the Schlenk line before reconstitution, which was carried out in an anaerobic chamber at 4°C in a cooling-heating block (Dry Bath H2O3-100C; Coyote Bioscience, Beijing, China). Dithiothreitol (DTT, 100 mM) was added to the purified protein fraction (100 μM) at a final concentration of 10 mM. The mixture was incubated for 45min before 2 equivalents of Fe $(NH_4)_2$ $(SO_4)_2$ solution (5 mM) were added every 10 min 4 times carefully (8 equivalents total). After incubation at 4 °C for 20 min, 2 equivalents of $Na_2S$ solution (20 mM) were added every 20 min for 4 times (8 equivalents total). The mixture was incubated overnight at 4°C and the color changed to brown. The excess iron and sulfide were removed by desalting on a PD-10 desalting column (GE) pre-equilibrated with desalting buffer (300 mM NaCl, 50 mM $NaH_2PO_4$, and 10% (v/v) glycerol at pH 8.0). The protein fractions were collected, concentrated and were used directly for in vitro assays or stored at -80 °C until further use. Protein concentration was determined by the Bradford assay.

**Analysis of iron and labile sulfide in ArsL**

The protocol for analyzing the content of Fe and labile sulfide was described previously. [2,3] To determine the Fe content, reagent A (equal volumes of 1.2 M HC1 and 4.5% (w/v) KMnO4) and reagent B (6.5 mM ferrozine, 13.1 mM neocuproine, 2 M ascorbic acid, 5 M ammonium acetate) were prepared in advance. One hundred microliters of reconstituted protein or iron standard at different concentrations was mixed with 50 μL of reagent A at 60 °C for 2 h. The mixture was then cooled and 10 μL of reagent B was added at room temperature for 30 min before the absorbance was recorded at 562 nm.

To determine the labile sulfide content, 100 μL of reconstituted protein (in 50 mM NaOH) or 200 μM $Na_2S$ solution (in 50 mM NaOH) at different concentrations was mixed with 300 μL of 1% (w/v) zinc acetate and 15 μL of 3 M NaOH and incubated for 2 h under anaerobic conditions. Subsequently, 75 μL of 0.1% (w/v) DMPD (in 5 M HCl) and 2 μL of 23 mM FeCl3 (in 1.2 M HCl) were added. The samples were removed from the anaerobic chamber and centrifuged at 14,000rpm for 15 min. Then, 600 μL of 2 mM EPPS (pH 8.0) was added to 400 μL of the supernatant from each standard and sample. The mixture was incubated at room temperature for 30 min before the absorbance was recorded at 666 nm.

**Expression and purification of ArsM.**

The codon-optimized gene fragment of ArsM (accession number WP_219608244) from *Burkholderia gladioli* GSRB05 (NZ_JAGSIB010000059.1)[1] was synthesized by GENE CREATE (Wuhan, China) and inserted into the vector pTrc-HisA (Amp). Using the Minerva Super Fusion Cloning Kit (UE EVERBRIGHT), the gene was cloned and inserted into the similarly digested expression vector pEt28a (+) (Kana). The primers used are listed in Table S1. The plasmid was subsequently used to transform the *E. coli* BL21 Δmtn (DE3) strain. The cells were grown in 1 liter LB medium at 37 °C and 180 rpm. When the $OD_{600}$ reached 0.8, the cultures were cooled down to 16 °C. Protein expression was induced by 0.3 mM IPTG. The cells were harvested after incubation at 16 °C for 18-20 h. The cell pellets from 2 liters culture were resuspended in 40 mL of MOPS buffer containing 50 mM MOPS, 500 mM NaCl, 5 mM imidazole and 10% (v/v) glycerol at pH 7.5. Then, 1 mM PMSF, 120 mg lysozyme and 2 mg DNase I were added, and the mixture was disrupted by sonication (Scientz, China) at 0 °C for 30 min (power 300 W, working 4 sec with a 6 sec interval). The cell debris was removed by centrifugation at 14,000g (Cence, China) for 45 min. The supernatant was incubated for 1 h with 3 mL of Ni-NTA resin (GenScript) preequilibrated with the MOPS buffer (50 mM MOPS, 500 mM NaCl, 5 mM imidazole and 10% (v/v) glycerol at pH 7.5). The Ni-NTA resin was loaded onto a polypropylene column and washed with 40 mL of MOPS buffer, followed by 300 mL of 30 mM imidazole in MOPS buffer. ArsM was eluted from the column with elution buffer (500 mM imidazole in MOPS buffer). The elution fractions were buffer-exchanged to 50 mM MOPS, 500 mM NaCl and 10% (v/v) glycerol at pH 7.5 using a 10-DG desalting column (GE Healthcare). The purified proteins were concentrated using Amicon Ultra 15 centrifugal filter devices (Millipore). Aerobically purified ArsM proteins were degassed by the Schlenk line before use.

**Ultraviolet–visible spectroscopy of ArsL.**

The UV-Vis spectra of ArsL (50 μM) were obtained on a UV-Vis spectrophotometer (Shimadzu, Japan) from 200 nm to 900 nm. The baseline was corrected with the desalting buffer used to prepare the sample. Under anaerobic condition, dithionite was added to the protein sample at a final concentration of 0.33 mM, and the mixture was incubated for 10 min at 4 °C, without dithionite as a control. These two samples were sealed in quartz cells (100 μL each) and removed from the glovebox for analysis. (Figure S2)

**Activity assay of the SAM cleavage reaction of ArsL.**

Under anaerobic condition, the reactions were assembled with 200 μM reconstituted ArsL, 2 mM SAM, 10 mM dithionite, 300 mM NaCl, 50 mM $NaH_2PO_4$, and 10% (v/v) glycerol at pH 8.0 in a final volume of 30 μL. The mixtures were incubated at 28 °C for 4 h. The control reactions without ArsL or dithionite were set up similarly by replacing the corresponding component with an equal volume of buffer. Then, the reactions were quenched with 30 μL of 10% TFA, followed by centrifugation to separate the precipitated proteins and the supernatant. The supernatant was analyzed by high-performance liquid chromatography (HPLC, Shimadzu) on a C18 column (5 μm, 4.6 mm × 150 mm, Shimadzu) monitored at 254 nm absorbance, using Method 1. Method 1: 3% solvent B for 3 min, a gradient from 3% to 19% solvent B for 3-13 min was followed by a gradient from 19% to 100% for 13-15 min and then 100% B for 15-18 min and 3% B for 18-24 min at a flow rate of 1 mL/min (solvent A: 0.1% aqueous TFA and solvent B: 0.1% TFA in acetonitrile).

To detect another product of the SAM cleavage reaction, a separate set of examples were derivatized with 2,4-dinitrofluorobenzene (DNFB) and analyzed by HPLC. After the reactions were quenched with 30 μL of 10% TFA and centrifuged, the supernatants were concentrated by a vacuum freeze dryer and redissolved in 30 μL of 1M $NaHCO_3$, 12 μL of acetonitrile and 1% DNFB solution in acetonitrile (3.5 μL). The reactions were heated at 40 °C for 3 h, cooled to rt and quenched by the addition of 2 N HCl (22.5 μL). The samples were analyzed by HPLC on a

C18 column (5 μm, 4.6 mm × 150 mm, Shimadzu) at an absorbance wavelength of 340 nm using Method 2. Method 2: 0% solvent B for 4 min and 19% solvent B for 4-7 min, a gradient from 19% to 24.5% solvent B for 7- 19 min; and a gradient from 24.5% to 45% solvent B for 15-36 min. Then a linear gradient of 45-100% B for 36-37 min, 100% solvent B for 37-39 min, and 0% solvent B for 39-43 min was used at a flow rate of 1 mL/min (solvent A: 0.1% aqueous TFA and solvent B: 0.1% TFA in acetonitrile).

**LC-MS analysis of the SAM cleavage reaction.**

Under anaerobic condition, the reactions were assembled with 150 μM reconstituted ArsL, 3 mM SAM, 10 mM dithionite, 300 mM NaCl, 50 mM $NaH_2PO_4$, and 10% (v/v) glycerol at pH 8.0 in a final volume of 100 μL. The mixture was incubated at 28 °C for 4 h. The control reaction omitting dithionite was also performed. Then, the reactions were quenched with 100 μL of 10% TFA, followed by centrifugation to separate the precipitated proteins and the supernatant. The supernatant was concentrated by a vacuum freeze dryer and redissolved in 50% methanol/$H_2O$. After centrifugation at 12,000 rpm for 30 min, 10 μL of the supernatant was subjected to liquid chromatography-high resolution mass spectrometry (LC-HRMS) analysis using a Q Exactive™ HF/UltiMate™ 3000 RSLCnano ( Thermo Fisher ) instrument. LC-MS analysis was carried out on a ZIC-HILIC column (5 mm, 200 Å, 150×4.6 mm; Merck). The HPLC procedure was as follows (Method 3): 90% solvent B for 2 min, a gradient from 90% to 70% solvent B for 2-5 min; a gradient from 70% to 50% solvent B for 5-30 min, and 90% solvent B for 30-40 min (solvent A: 90% 0.1 M ammonium acetate and 10% acetonitrile; solvent B: acetonitrile). The flow rate was set to 0.75 mL/min. The mass spectrometer was run in ESI positive mode.

**LC-MS analysis of the purified product.**

The HPLC fractions for MS were concentrated by a vacuum freeze dryer and redissolved in water. A 10μL portion of the supernatant was analyzed by a miorOTOF QII LC/MS instrument (Bruker Daltonics Inc., USA) on a C18 column (3 μm, 2.1 mm×100 mm; Shimadzu). The HPLC conditions were as follows (Method 4): a linear gradient of 5-8% solvent A for 0-7 min, a linear gradient of 8-95% solvent A for 7-12min, 95% B for 12-19 min and 5% A for 19-32 min at a flow rate of 0.2 mL/min. (solvent A: 0.1% formic acid in acetonitrile and solvent B: 0.1% aqueous formic acid). The mass spectrometer was run in ESI positive or negative mode.

**Analysis of reaction products.**

Under anaerobic condition, the reactions were assembled with 200 μM reconstituted ArsL, 2.5mM SAM, 10 mM dithionite, 2 mM As (III), 300 mM NaCl, 50 mM $NaH_2PO_4$, and 10% (v/v) glycerol at pH 8.0 in a final volume of 50 μL. The mixtures were incubated at 28°C for 4 h. The control reactions without ArsL or dithionite were set up similarly by replacing the corresponding component with an equal volume of buffer. Then, 12 μL of the reaction mixture was quenched with 12 μL of 10% TFA, followed by centrifugation to separate the precipitated proteins and the supernatant. The supernatant was analyzed by HPLC on a C18 column (5 μm, 4.6mm × 150 mm; Shimadzu) monitored at 254 nm absorbance, using Method 1.

The products of ArsL were detected by derivatization with L-FDAA (Marfey's Reagent)[4]. The remaining reaction mixture (38μL) was treated with 6% (v/v) $H_2O_2$ at room temperature for

10min. Then, the mixture was quenched with 47.5 μL of 10% TFA. The supernatant was obtained by centrifugation, concentrated by a vacuum freeze dryer, and redissolved in 38 μL of 1 M $NaHCO_3$, 20 μL of acetonitrile, and 1% L-FDAA solution in acetone (5.2 μL). The reactions were heated at 50°C for 1 h, cooled down to room temperature and quenched by the addition of 2 N HCl (30 μL). Standard amino acid (2-aminobutyric acid, ABA) was reacted with L-FDAA in the same way (Figure S5). The sample was analyzed by HPLC on a C18 column (5 μm, 4.6 mm × 150 mm, Shimadzu) at an absorbance wavelength of 340 nm using Method 2.

**Different reductants for ArsL.**

Under anaerobic condition, reactions were assembled with 200 μM reconstituted ArsL, 2.5 mM SAM, 2 mM As (III), no reductant/with 10 mM dithionite/with 5 mM Ti (III) citrate/with 1mM methyl viologen (MV) and 4 mM NADPH, 300 mM NaCl, 50 mM $NaH_2PO_4$, and 10% (v/v) glycerol at pH 8.0, in a final volume of 50 μL. The mixtures were incubated at 28°C for 4 h. The control reaction without ArsL was set up similarly by replacing the corresponding component with an equal volume of buffer. The analysis of enzyme activity and products was carried out as described in the analysis of reaction products.

**Coupled assay of ArsL and ArsM.**

Under anaerobic condition, reactions were assembled with 200 μM reconstituted ArsL, 200 μM ArsM, 3.5 mM SAM, 5 mM Ti (III) citrate, 2 mM As (III), 300 mM NaCl, 50 mM $NaH_2PO_4$, and 10% (v/v) glycerol at pH 8.0 in a final volume of 65 μL. A control reaction without ArsM was similarly prepared by replacing the corresponding component with an equal volume of buffer. The mixture was incubated at 28°C for 5 h. Then, 15 μL of reaction mixture was quenched with 15 μL of 10% TFA, followed by centrifugation to separate the precipitated proteins and the supernatant. The supernatant was analyzed by HPLC on a C18 column (5 μm, 4.6mm × 150 mm; Shimadzu) monitored at 254 nm absorbance using Method 1. The remaining reaction mixture (50μL) was treated with 6% (v/v) $H_2O_2$ at room temperature for 10 min. The products of the coupled reaction of ArsL and ArsM were detected by derivatization with L-FDAA.

**Direct detection of AST-OH and AST by LC-MS.**

The ArsL reaction mixture (200 μL) containing 200 μM reconstituted ArsL, 3 mM SAM, 5 mM Ti (III) citrate, 3 mM As (III), 300 mM NaCl, 50 mM $NaH_2PO_4$, and 10% (v/v) glycerol at pH 8.0 was incubated at 28°C for 5 h under anaerobic condition. The control reaction without reductant was similarly prepared by replacing the corresponding component with an equal volume of buffer. And the coupled reaction of ArsL and ArsM mixture (200μL), containing 200 μM reconstituted ArsL, 200 μM ArsM, 3 mM SAM, 5 mM Ti (III) citrate, 3 mM As (III), 300 mM NaCl, 50 mM $NaH_2PO_4$, and 10% (v/v) glycerol at pH 8.0, was also incubated at 28°C for 5 h under anaerobic condition. The reactions were oxidized by 6% (v/v) $H_2O_2$ at room temperature for 10 min, followed by the addition of 250 μL of 10% TFA, after which the supernatant was separated by centrifugation. The supernatant was concentrated by a vacuum freeze dryer and redissolved in 50% methanol/$H_2O$. After centrifugation, 10 μL of the supernatant was subjected to LC-HRMS analysis using a Q Exactive™ HF/UltiMate™ 3000 RSLCnano (Thermo Fisher) instrument. LC-MS analysis was carried out on a ZIC-HILIC column (5 mm, 200 Å, 150×4.6 mm; Merck) using Method 3.

**Stepwise cascade reaction of ArsL and ArsM**

The ArsL reaction mixture (200 µL) containing 240 µM reconstituted ArsL, 3 mM SAM, 5 mM Ti (III) citrate, 3 mM As (III), 300 mM NaCl, 50 mM $NaH_2PO_4$, and 10% (v/v) glycerol at pH 8.0 was incubated at 28°C for 4 h under anaerobic condition. After that, the reaction was quenched by heating at 50°C for 10min, followed by centrifugation to separate the precipitated proteins and the supernatant. Half of the supernatant was oxidized by 6% (v/v) $H_2O_2$ at room temperature for 10 min (reaction 2), and the other half was left untreated (reaction 1). The supernatants were cooled under anaerobic condition and concentrated by a vacuum freeze dryer. The reaction vials were transferred back to an anaerobic chamber, redissolved in 55 µL of $H_2O$ and adjusted to pH 7.5 with 0.5 M NaOH. Add 2 mM SAM, 240 µM ArsM, 500 mM NaCl, 50 mM MOPS, and 10% (v/v) glycerol at pH 7.5 to the reaction vials. The solutions were incubated at 37°C for 8 h under anaerobic condition. After that, the reactions were treated with 6% (v/v) $H_2O_2$ at room temperature for 10 min and quenched with an equal volume of 10% TFA. The supernatant was obtained by centrifugation and was subsequently, concentrated by a vacuum freeze dryer. The products were detected by derivatization with L-FDAA (Figure S6).

**Activity assay of ArsL$_{\Delta C}$.**

Under anaerobic condition, the reactions were assembled with 200 µM reconstituted wild-type ArsL or ArsL$_{\Delta C}$, 2.5mM SAM, 2 mM As (III), 5 mM Ti (III) citrate, 300 mM NaCl, 50 mM $NaH_2PO_4$, and 10% (v/v) glycerol at pH 8.0 in a final volume of 50 µL. The mixtures were incubated at 28°C for 4h. The control reactions without As (III), enzyme or Ti (III) citrate were set up similarly by replacing the corresponding component with an equal volume of buffer. The analysis of enzyme activity and products was carried out as described in the analysis of reaction products.

**Different substrates for ArsL.**

Under anaerobic condition, the reactions were assembled with 250 µM reconstituted wild-type ArsL or 230 µM reconstituted ArsL$_{\Delta C}$**,** 2 mM SAM, 5 mM Ti (III) citrate, no substrate/ 10 mM phosphorous acid ( $H_3PO_3$ ) / $C_2H_7OP$ / $C_3H_9P$ , 300 mM NaCl, 50 mM $NaH_2PO_4$, and 10% (v/v) glycerol at pH 8.0 in a final volume of 50 µL. The mixtures were incubated at 28°C for 4 h. The control reactions without ArsL or dithionite were set up similarly by replacing the corresponding component with an equal volume of buffer. Then, 12 µL of the reaction mixture was quenched with 12 µL of 10% TFA, followed by centrifugation to separate the precipitated proteins and the supernatant. The supernatant was analyzed by HPLC on a C18 column (5 µm, 4.6 mm × 150 mm; Shimadzu) at an absorbance wavelength of 254 nm using Method 1.

The products of wild-type ArsL or ArsL$_{\Delta C}$ in the presence of $H_3PO_3$, $C_2H_7OP$, or $C_3H_9P$ were detected by derivatization with L-FDAA (Marfey's Reagent)[4]. The mixture was quenched with 38 µL of 10% TFA. The supernatant was obtained by centrifugation, concentrated by a vacuum freeze dryer, and redissolved in 38 µL of 1 M NaHCO3, 19 µL of acetonitrile, and 1% L-FDAA solution in acetone (5 µL). The reactions were heated at 40°C for 3 h, cooled down to room temperature and quenched by the addition of 2 N HCl (31 µL). The sample was analyzed by HPLC on a C18 column (5 µm, 4.6 mm × 150 mm, Shimadzu) at an absorbance wavelength of 340 nm using Method 2.

**Table S1: List of primers used in this study**

| Primer name | | Sequence (5’-3’) |
|---|---|---|
| ArsL-1 | Forward | GGCTAGCGGCGGCGGCGGCGGCGGCATGGCCAATTATCTGG TGGTTAGTACCTTTG |
| | Reverse | TTAGTCTGACCATCTCATCTGTAACATCATTGGC |
| ArsL-2 | Forward | GGCAAAGGTAGCGTTGCCAATGATGTTACA |
| | Reverse | AAGCGTGGCGCTTTCTCATAGCTC |
| ArsL-3 | Forward | CTGAGATACCTACAGCGTGAGCTATGAG |
| | Reverse | GCCATGCCGCCGCCGCCGCCGCCGCTAGCCATACCATGATGA TGATGATGATGAGAAC |
| ArsM-1 | Forward | TCGCGGATCCATGGAAATGGATAGTGTGATTC |
| | Reverse | GCAGCTGTTGTAGCGCAGATTGACTCGAGCACC |
| ArsM-2 | Forward | CGCAGATTGACTCGAGCACCACCACCACCACCACTGAGAT |
| | Reverse | ACTCAAAGGCGGTAATACGGTTATCCACAG |
| ArsM-3 | Forward | TCTGTGGATAACCGTATTACCGCCTTTGAG |
| | Reverse | CCATTTCCATGGATCCGCGACCCATTTGCTGTCCAC |
| ArsL$_{\Delta C}$-1 | Forward | TTTTTAAACGCTGAGAGAAGATTTTCAGCCTGATACAGATTA AATCAG |
| | Reverse | GTTGGTAGCTCTTGATCCGGCAAAC |
| ArsL$_{\Delta C}$-2 | Forward | GTTTGCCGGATCAAGAGCTACCAAC |
| | Reverse | GTATCGTCGTATCCCACTACCGAGATATC |
| ArsL$_{\Delta C}$-3 | Forward | ATATCTCGGTAGTGGGATACGACGATAC |
| | Reverse | GCTGAAAATCTTCTCTCAGCGTTTAAAAACACCCGGTTCCGG |

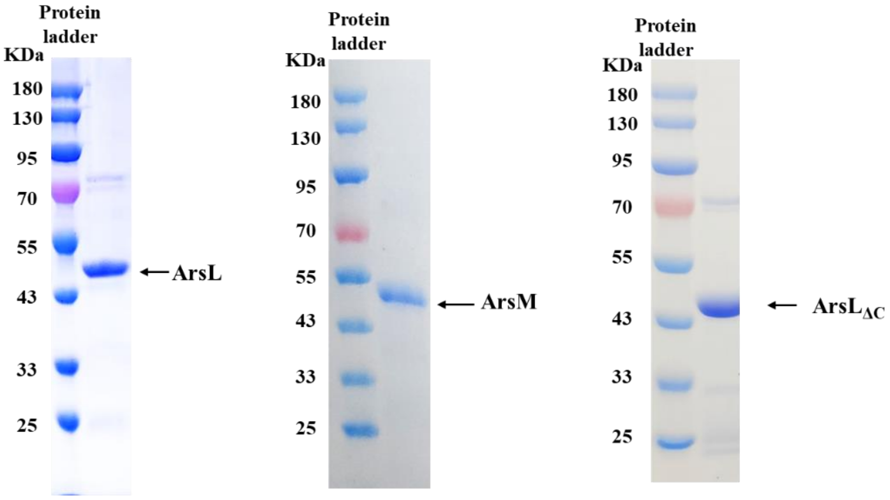


**Figure S1**. Coomassie blue stained SDS-PAGE gel showing purified ArsL, ArsM, and ArsL$_{\Delta C}$ proteins.

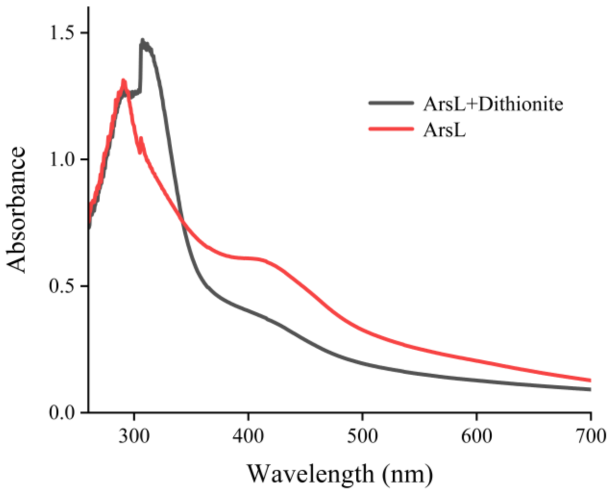


**Figure S2**. Ultraviolet-visible absorption spectra of anaerobically isolated (red) and dithionite-reduced (black) ArsL.

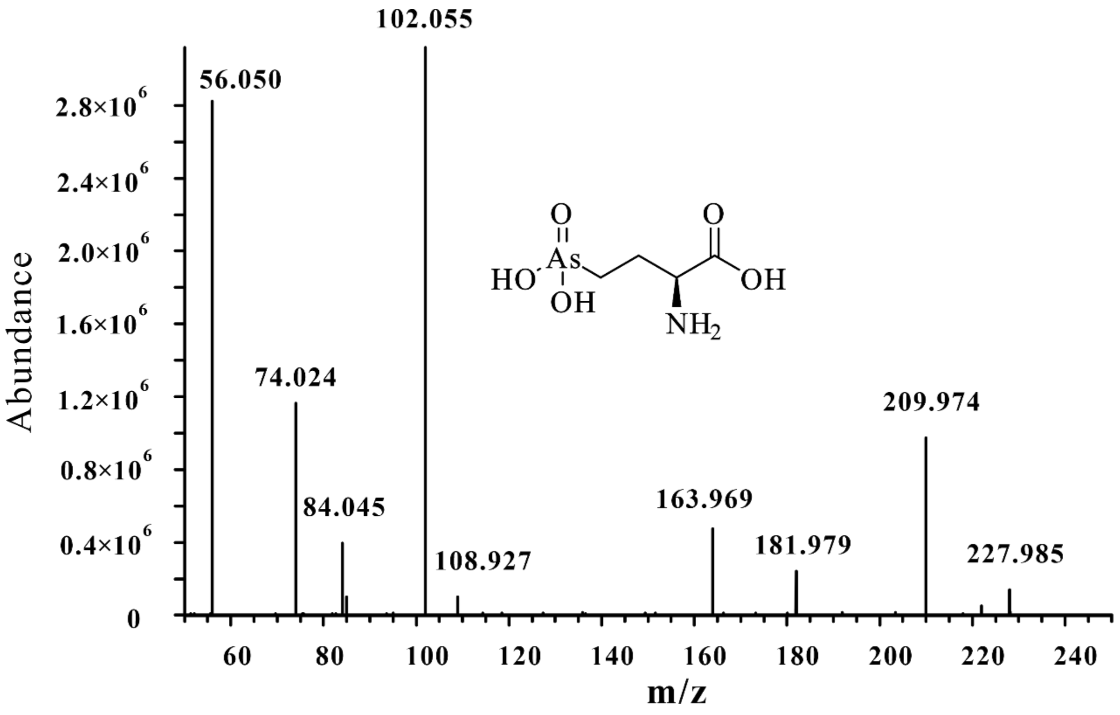


**Figure S3**. HR-MS/MS fragmentation of 2-amino-4-(dihydroxyarsonoyl)butanoic acid (AST-OH) in the ArsL reaction. The MS/MS fragmentation of AST-OH was identical to the literature, [5] and included m/z 55.6 ($[C_3H_6N]^+$), 73.7 ($[C_3H_8ON]^+$), 102.2 ($[C_4H_8O_2N]^+$), 164.6 ($[C_2H_3O_3NAs]^+$), 181 ($[C_2H_5O_4NAs]^+$), and 209.9 ($[C_4H_9O_4NAs]^+$).

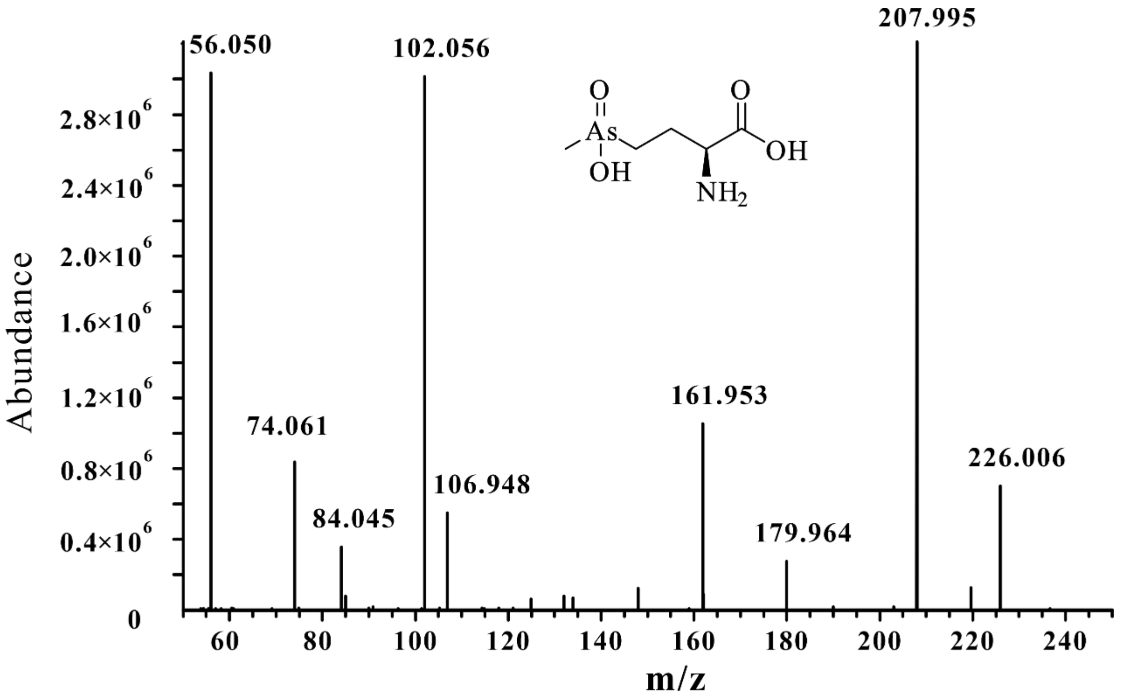


**Figure S4**. HR-MS/MS fragmentation of arsinothricin (AST) in the cascade reaction of ArsL with ArsM.. The MS/MS fragmentation of AST-OH was identical to the literature [5], and included m/z 56.0502($[C_3H_6N]^+$), 74.0606 ($[C_3H_8ON]^+$), 102.0553 ($[C_4H_8O_2N]^+$),106.9476 ($[CH4OAs]^+$), 161.9531 ($[C_3H_5O_2NAs]^+$), 179.9636 ($[C_3H_7O_3NAs]^+$) and 207.9950 ($[C_5H_{11}O_3NAs]^+$).

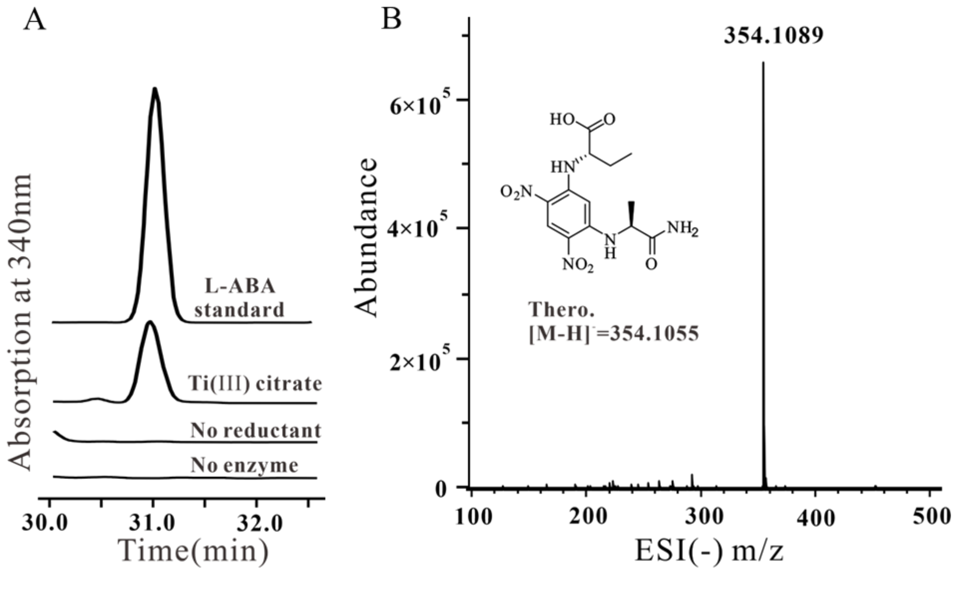


**Figure S5**. Detection of L-ABA from the ArsL reactions with Ti (III) citrate. A) HPLC elution profiles of the L-FDAA derivatives of L-ABA. B) Negative ionization high-resolution mass spectrum of the L-FDAA derivatives of the L-ABA standard.

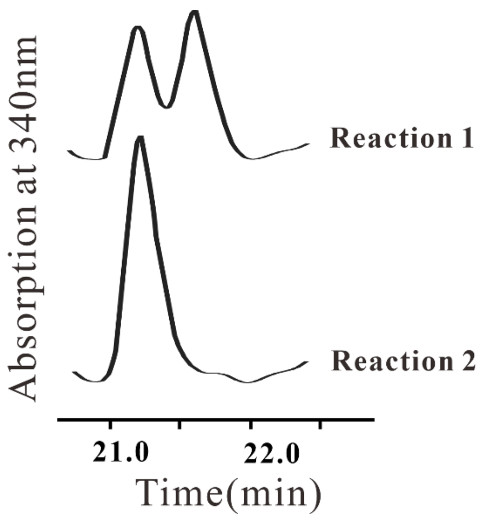


**Figure S6**. HPLC elution profiles of the L-FDAA derivatives of AST-OH and AST from the different reaction of ArsL with ArsM. Reaction 1: reaction with the unoxidized product of ArsL and ArsM; Reaction 2: reaction with the oxidized product of ArsL and ArsM.

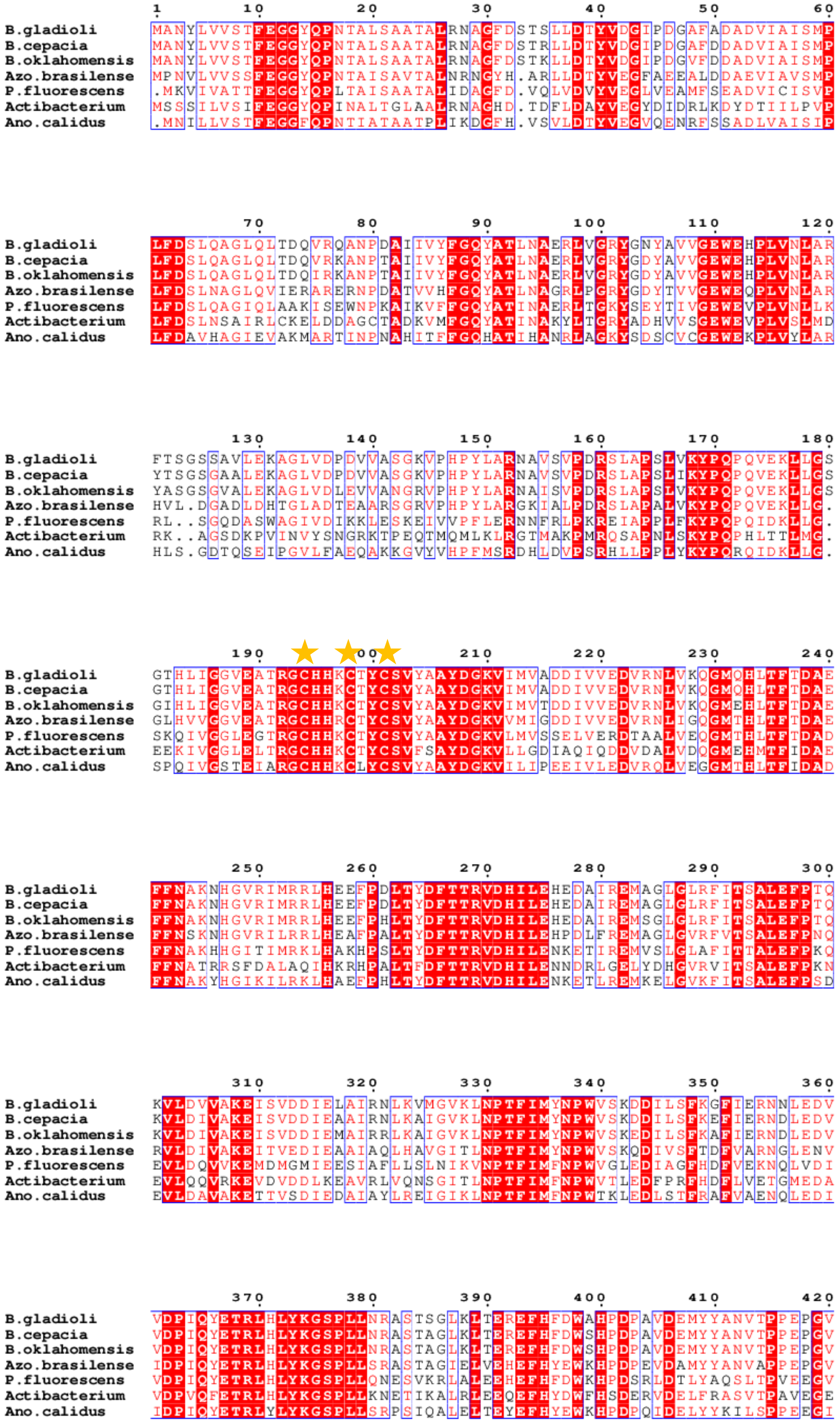
B.gladioli
B.cepacia
B.oklahomensis
Azo.brasilense
P.fluorescens
Actibacterium
Ano.calidus

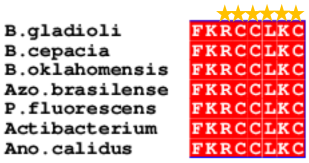


**Figure S7**. MEGA 11 and ESPript 3.0 multiple alignments of *Bg*ArsL and its homologs. The protein sequence of *Bg*ArsL (accession number WP_219608243) was compared with orthologs from *Burkholderia cepacia* (WP_060174813.1), *Burkholderia oklahomensis* (accession number WP_038802160.1), *Azospirillum brasilense* (accession number WP_211101616.1), *Pseudomonas fluorescens* (accession number WP_214918812.1), *Actibacterium sp.* 188UL27-1 (accession number WP_204414396.1) and *Anoxybacillus calidus* (accession number WP_181538902.1).

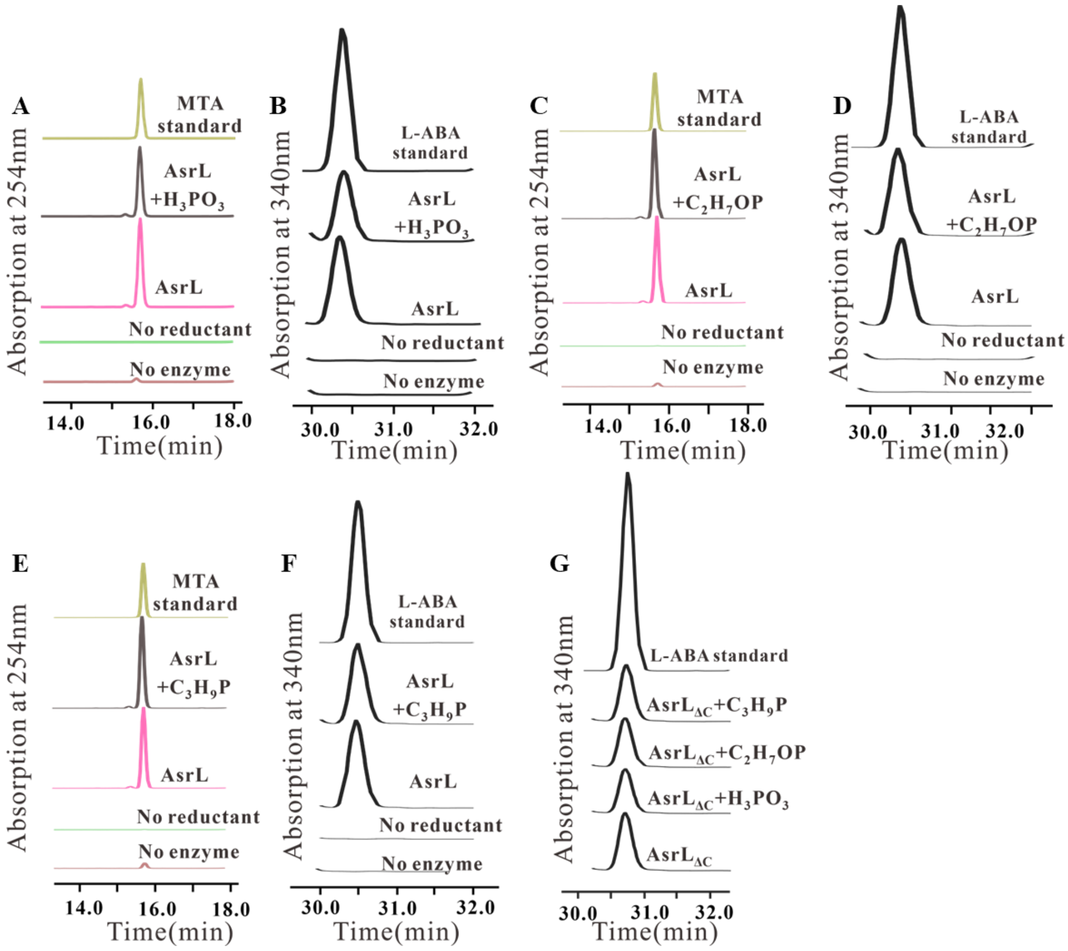


**Figure S8**. Detection of product from the ArsL or $ArsL_{\Delta C}$ reactions in the presence of $H_3PO_3$, $C_2H_7OP$, $C_3H_9P$. A) HPLC traces of the SAM cleavage reactions by ArsL in the presence of $H_3PO_3$. B) HPLC elution profiles of the L-FDAA derivatives by ArsL in the presence of $H_3PO_3$. C) HPLC traces of the SAM cleavage reactions by ArsL in the presence of $C_2H_7OP$. D) HPLC elution profiles of the L-FDAA derivatives by ArsL in the presence of $C_2H_7OP$. E) HPLC traces of the SAM cleavage reactions by ArsL in the presence of $C_3H_9P$. F) HPLC elution profiles of the L-FDAA derivatives by ArsL in the presence of $C_3H_9P$. G) HPLC elution profiles of the L-FDAA derivatives by $ArsL_{\Delta C}$ in the presence of $H_3PO_3$, $C_2H_7OP$, or $C_3H_9P$.

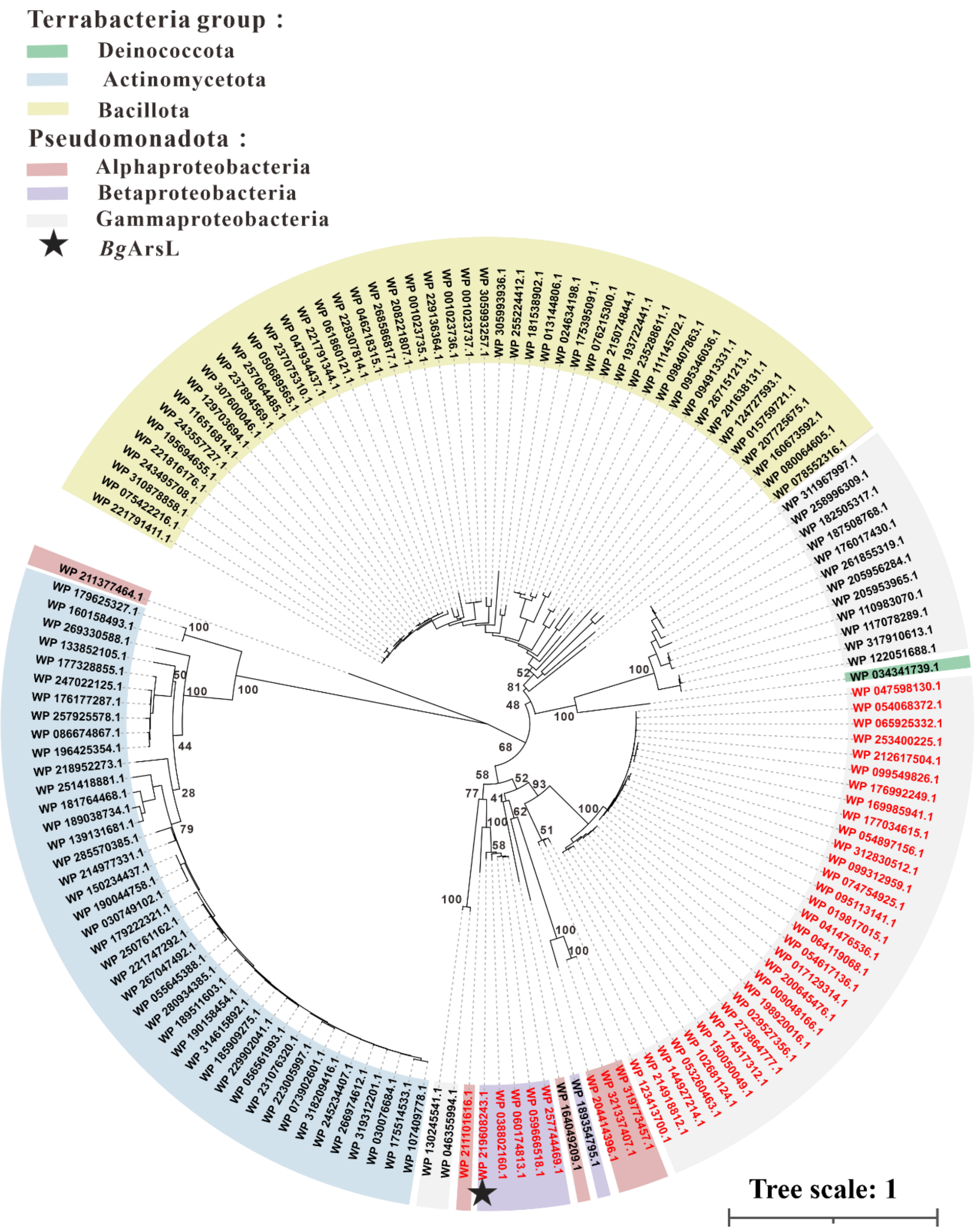


**Figure S9**. Phylogenetic analysis of *Bg*ArsL homologs in the RCCLKC-tail radical SAM protein (NCBI HMM accession NF040542.1). The phylogenetic tree was constructed using MEGA11 software. The accession numbers of radical SAM protein ArsL are indicated in red.
